# Temporal Ablation of the Ciliary Protein IFT88 Alters Normal Brainwave Patterns

**DOI:** 10.1101/2024.04.03.587983

**Authors:** Matthew R. Strobel, Yuxin Zhou, Liyan Qiu, Aldebaran M. Hofer, Xuanmao Chen

## Abstract

The primary cilium is a hair-like organelle that hosts molecular machinery for various developmental and homeostatic signaling pathways. Its alteration can cause rare ciliopathies such as the Bardet-Biedl and Joubert syndromes, but is also linked to Alzheimer’s disease, clinical depression, and autism spectrum disorder. These afflictions are caused by disturbances in a wide variety of genes but a common phenotype amongst them is cognitive impairment. While cilia-mediated neural function has been widely examined in early neurodevelopment, their function in the adult brain is not well understood. To help elucidate the role of cilia in neural activity, we temporally induced the ablation of IFT88, a gene encoding the intraflagellar transport 88 protein which is neccessary for ciliogenesis, in adult mice before performing memory-related behavioral assays and electroencephalogram/electromyogram (EEG/EMG) recordings. Inducible IFT88 KO mice exhibited severe learning deficits in trace fear conditioning and Morris water maze tests. They had strongly affected brainwave activity both under isoflurane induced anesthesia and during normal activity. And additionally, inducible IFT88 KO mice had altered sleep architecture and attenuated phase-amplitude coupling, a process that underlies learning and memory formation. These results highlight the growing significance of primary cilia for healthy neural function in the adult brain.

## Introduction

Primary cilia are specialized organelles that emanate from the plasma membrane of nearly all vertebrate cells. And despite being contiguous with the cellular membrane, the ciliary membrane maintains a unique lipid and protein composition ^1,2^. The cilium also has a filter-like transition zone at the basal end which requires chaperoning and an intraflagellar transport system to traverse for proteins with a stokes radii > 8nm ^3,4^. This membrane composition and barrier to diffusion isolates the cilium from the cell body and allows it to operate as its own unique microdomain.

Numerous intracellular signaling pathways have been linked to primary cilia including Sonic Hedgehog (Shh), WNT, Notch, HIPPO, PDGF, mTOR, and G-protein coupled receptor (GPCR)-mediated cAMP pathways. Primary cilia therefore regulate many key developmental, sensory, and homeostatic signals ^1,5–7^. Mutations to genes in these pathways or other ciliary genes can lead to severe developmental disorders known as ciliopathies. To date over 60 ciliary genes have been found to impair neurological development leading to conditions such as Joubert, Meckel, and Bardet-Biedl syndromes ^8^. Outside of these classically identified ciliopathies, ciliary genes have also been connected to other neurological conditions. The autism spectrum disorder genes FXS and AHI1 have been linked through genome-wide association studies to Joubert Syndrome and may act as signal transducers of Shh ^9–11^. Disruption of these genes leads to reduced neuronal differentiation, dendritic complexity, memory deficits, and reduced ciliation of neurons in the hippocampal dentate gyrus ^12–14^. In Alzheimer’s disease (AD) the ciliary receptors p75^NTR^, SSTR3, and 5HTR6 have been implicated in the progression of cognitive decline. And it has been shown that p75^NTR^ can be aberrantly activated by amyloid-beta (Aβ) peptides resulting in tyrosine kinase mediated apoptosis in the hippocampus ^15–17^.

Building on its role in neurological disease, impaired cognition and memory have been recorded in other ciliary studies. Aβ peptides can bind and activate the Gα_i_ linked ciliary GPCR SSTR3, reducing cAMP levels and long-term potentiation (LTP), resulting in impaired novel object recognition and spatial recall ^18,19^. Chemical and genetic manipulations of 5HTR6 in an Alzheimer’s-like APP/PS1 mouse model modulates both ciliary length and cognitive ability^20^. Additionally, hippocampal ablation of IFT88, a transport protein necessary for cilia formation, impairs distant memory formation. Finaly, forebrain-specific IFT88 KO mice struggle with novel object recognition, Morris water maze performance, and aversive fear conditioning, possibly through destabilization of perineuronal nets ^21,22^.

One ciliary signaling protein that may give insight to how the cilium is linked to neural function is type 3 adenylyl cyclase (AC3). Originally identified in olfactory cilia, AC3 generates cAMP to mediate the activation of CNG ion channels resulting in rapid depolarization of olfactory neurons ^23^. AC3 in the central nervous system is not well studied but it has become a protein of interest after human genome-wide association studies found a strong relation between AC3 and major depressive disorder (MDD) ^24,25^. AC3 is well known as a protein marker of the neuronal primary cilium. In mice ablation of AC3 reduces neuronal activity, and causes various depression-like phenotypes, including altered sleep patterns, similar to those seen in human patients ^26^.

While all of these findings indicate a strong link between primary cilia and neural function, the use of behavioral paradigms presents its own set of challenges due to varying interpretations of test results, qualitative scoring, and behavioral variability. To quantitatively examine if cilia influence sleep and brainwave patterns we performed 24-hour continuous 3-channel EEG/EMG recordings on inducible global IFT88 knockout (KO) mice (IFT88 flox/flox UBC-Cre/ERT2, tamoxifen-inducible), permitting temporal ablation of primary cilia. We also subjected these mice to behavioral paradigms to assess how memory formation was affected in our model.

Brain EEG waveforms were analyzed to assess if cilia deletion alters basal neuronal activity. Cross-frequency-coupling (CFC) between theta and gamma frequency oscillations was calculated to better understand the cognitive impairments that have been seen in various ciliary ablation models, since CFC is known to participate in synaptic plasticity and consolidating short-term memory into long-term memory ^27–31^. Our results show that temporal ablation of IFT88 impairs memory formation, alters sleep patterns, reduces EEG waveform activity, and attenuates theta-gamma phase-amplitude coupling (PAC). These results support the notion that primary cilia are necessary for routine neural function. Additionally, our results are set apart from other studies in the field by expanding on our behavioral assays with quantitative EEG analysis of brain activity, power, and synchronization. This is the first study to show that loss of primary cilia influences PAC, which helps elucidate the role neuronal cilia play in the synaptic functions which underlie memory formation and persistence.

## Results

### Inducible IFT88 KO mice exhibit severe deficits in spatial learning and trace fear conditioning, and have reduced EEG activity under isoflurane-induced anesthesia

To study the role primary cilia play in neurological function we selected an inducible IFT88 knockout mouse line which causes structural extirpation of primary cilia ensuring disruption of all ciliary specific signaling (IFT88 flox/flox; UBC-Cre/ERT2 mice)^32,33^. In contrast, the neuronal AC3 knockout line does not alter primary cilia structure and is expected mainly to impair the ciliary cAMP signaling cascade downstream of AC3. Total abolishment of ciliary signaling pathways through IFT88 deleteion ensures a maximal effect which may be necessary to discern subtle alterations in the EEG signal. Although this model is not tissue-specific, it mimics the potential effects of a loss-of-function mutation in an adult organism.

There are complications with this IFT88 line however they can be minimized through proper experimental design. IFT88 has extraciliary roles that aid in cell division and migration, necessitating its expression during early development^36^. Although we cannot entirely exclude that residual cytosolic IFT88 may influence experimental results, the inducible nature of our model ensures that IFT88 protein and primary cilia are deleted in adult mice after the brain has already developed, thereby excluding neurodevelopmental complications. Conditional IFT88 silencing can also lead to cystic kidneys however these mice were shown to remain healthy with no signs of kidney disease two months after treatment and do not have significant kidney deficits or other outwardly apparent phenotypes until nine months after IFT88 deletion ^37^. To reduce the risk of health complications caused by conditional IFT88 silencing all EEG recordings were performed six weeks after the completion of tamoxifen treatment. Additionally, a global KO rather than a targeted KO was used to ensure a strong effect can be accurately separated from signal noise in the 2-channel EEG recording.

Six weeks after tamoxifen treatment we confirmed >90% cilia ablation (**Figure 1a**) and subjected KO and WT mice to several behavioral paradigms to characterize if hippocampal memory was impacted in our mouse line. First, for the fear conditioning test mice were trained to associate an innocuous tone with a mild but aversive electrical foot shock. During training the control animals had a markedly increased freezing score starting from the 3rd or 4th training cycle, consistent with what has been previously reported, indicating their ability to associate the aural tone with foot shock^34^. In contrast, inducible IFT88 KOs displayed a delayed learning response and significantly reduced freezing behaviors during the trace period of the seven training cycles (**Figure 1b**). To evaluate working memory persistence, we compared mouse freezing behavior from 10 minutes after training relative to 10 minutes prior to training, as short-term aversive memory lingers for several minutes. Trace fear conditioning significantly increased the relative freezing ratio of control mice, but there was no change in the inducible IFT88 KO (**Figure 1c**). Additionaly, In recall testing performed 2-3 hours after training KO mice displayed weaker tone-dependent freezing (**Figure 1d**). These data indicate that the inducible IFT88 KO mice exhibit marked deficits in associative learning.

**Figure 1:**
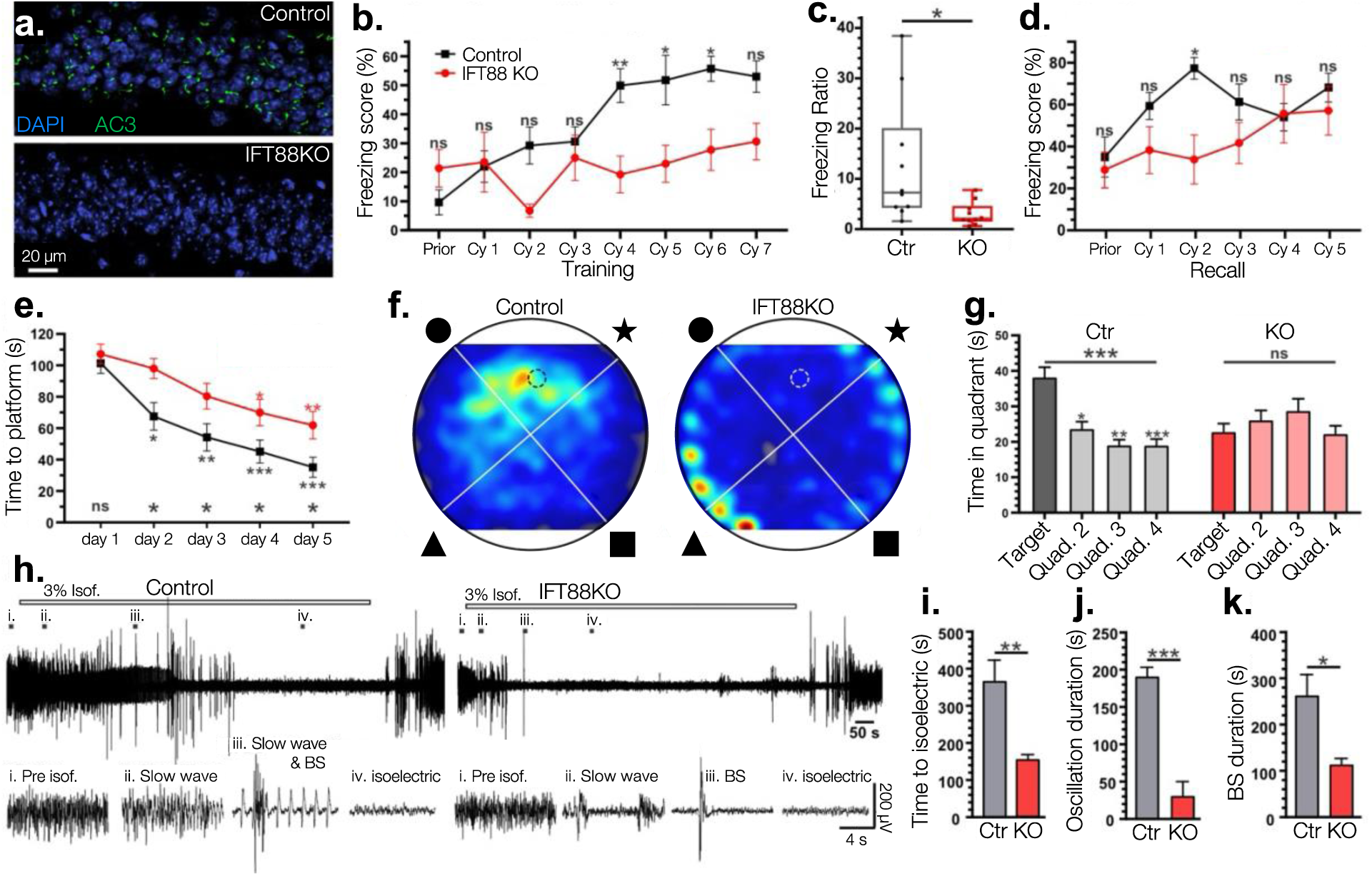
Inducible IFT88 KO mice exhibit severe learning deficits in hippocampal memory dependent paradigms and decreased EEG activity under anesthesia. (**a.**) Immunofluorescence staining using anti-AC3 antibody eight weeks post tamoxifen treatment. Efficiency of the inducible gene ablation was calculated by the ratio of cilia relative to DAPI cell counts in the CA1 region. The ciliation of WT controls was 73.7% ± 6.7%, n = 2 mice (4 sections), whereas KO ciliation was only 1.7% ± 1.4%, n = 2 mice (6 sections). (**b.**) In the trace fear conditioning test KO mice did not learn to associate the aural tone with a foot shock. After four training cycles they showed significant learning impairment compared to WT control mice. Bonferroni’s multiple comparisons test of two-way ANOVA, controls vs KO mice; n = 10 mixed sex littermates. (**c.**) When the freezing score from 10 minutes prior to training was ratioed against the freezing score from 10 minutes post training KO mice had a significantly lower ratio than WT mice, indicating significant learning deficiency; unpaired Student’s T-test. (**d.**) Freezing scores during recall testing showed a slight but significant reduction in KO mice. Bonferroni’s multiple comparisons test of two-way ANOVA, controls vs KOs. (**e.**) In the Morris water maze test control mice memorized the location of the hidden platform more quickly than IFT88 KO mice. Repeated measure one-way with post hoc Tukey’s multiple comparisons test between training day 1 and every other day. Bonferroni’s multiple comparisons test of two-way ANOVA, n = 10 mixed sex littermates (**f.**) A representative heatmap of time spent searching for the hidden platform where color intensity represents more time spent in the location. (**g.**) Whereas IFT88 mice spent a similar amount of time in each quadrant of the water maze, control mice spent significantly more time in the target quadrant which contained the hidden platform. Repeated measure one-way with post hoc Tukey’s multiple comparisons test. (**h-k**) Inducible IFT88 KOs require less time than controls to be fully anesthetized and reach the isoelectric stage (brain death). (**h.**) EEG recording during isoflurane-induced anesthesia. 3% isoflurane (Isof.) was used to induce anesthesia. Left: Control; Right: Inducible IFT88 KO. EEG traces at different anesthesia stages are enlarged below. (i), prior to isoflurane anesthesia (awake); (ii-iii), slow wave oscillation and burst suppression (BS) period; (iv), the isoelectric brain-death state. (**i-k**) IFT88 KO mice require less time to reach the isoelectric state, have reduced slow oscillations, and a shortened period of burst suppression. Anesthetic EEG data were collected from 12 WT and 9 KO mice (mixed sexes). Unpaired Student T-test between WT and KO mice. ns, not significant; *, p<0.05; **, p<0.01; ***, p<0.001.

Next, we subjected mice to the Morris water maze test to evaluate their ability to form spatial memories. In the Morris water maze test all mice had five days of training in which a hidden platform was placed at a fixed position, with four shapes attached around the chamber as spatial cues. If mice could not locate the platform after 120s they were placed on the platform to be shown its position. During these training days control mice showed significantly decreasing escape latency with each day as they became familiar with the hidden platform location, whereas IFT88 KO mice had a delayed ability to locate the platform (**Figure 1e**). In the subsequent probe test where the hidden platform was removed, control mice demonstrated a high preference for the target quadrant that previously contained the platform than any other quarters. But KO mice were unable to distinguish between the four quadrants and spent an equal amount of time in each (**Figure 1f-g**). These data illustrate that inducible IFT88 ablation impairs the formation of spatial memories and hampers navigational ability.

To monitor if general electrical activity in inducible IFT88 KO mice was suppressed we induced the isoelectric (brain-death) anesthetic state using 3% isoflurane while recording EEG activity. First, we found that IFT88 KO mice exhibited reduced EEG waveform activity with significantly lower electrical amplitude. (**Figure 1h**). Second, inducible IFT88 KOs required less time under isoflurane to reach the isoelectric state (**Figure 1i**). Furthermore, the duration of slow wave oscillation (**Figure 1j**), the spike number during slow-wave (frequency < 1 Hz), bursting numbers (**Figure 1h**), and duration of burst suppression (**Figure 1k**) under isoflurane-induced anesthesia were drastically decreased in KOs compared to littermate controls. This initial EEG experiment shows that brain waveform patterns in inducible IFT88 KOs that lack primary cilia are drastically altered, an interesting finding that warranted additional investigation as seen in the experiments below.

### Cilia ablation alters sleep bout patterns

To further examine how brainwave patterns are perturbed in inducible IFT88 KO mice, we conducted EEG analysis during sleep and awake cycles. Sleep disturbances including light sleep, shifts in sleep architecture, and increased rapid eye movement (REM) cycles are common phenotypes of major depressive disorder and have been observed in a transgenic AC3 knockdown mouse model for depression ^26,35^.

To observe if IFT88 ablated mice have similar sleep phenotypes to AC3 ablated mice, 24-hour EEG/EMG recordings were acquired, manually scored, and analyzed for alterations in sleep patterning. Typical trace recordings illustrating the method for scoring are shown in **Supplementary Figure 1**. On a percentage basis, inducible IFT88 KO mice spent comparable time awake, in non-REM sleep, or in REM sleep as their wild type littermates (**Figure 2a**). However, the average length of their non-REM sleep bouts and wake bouts increased significantly and almost doubled in length (p=0.0454 and p=0.0179; **Figure 2b**). In addition, these mice were also significantly less likely to be aroused from sleep, showing an 82% decrease in transitions from non-REM sleep to awake state (p=0.0024) and an 86% decrease in arousals from REM sleep when in the light cycle (**Figure 2c**). These observations also carried over into the dark cycle, traditionally when mice spend much less time sleeping, with these mice showing an 80% decrease in arousals from non-REM sleep (p=0.0459). These results differ slightly from those reported for AC3 KO mice, likely attributed to the different signaling pathways that are being disrupted as AC3 deletion does not structurally ablate primary cilia, whereas IFT88 deletion does.

**Figure 2:**
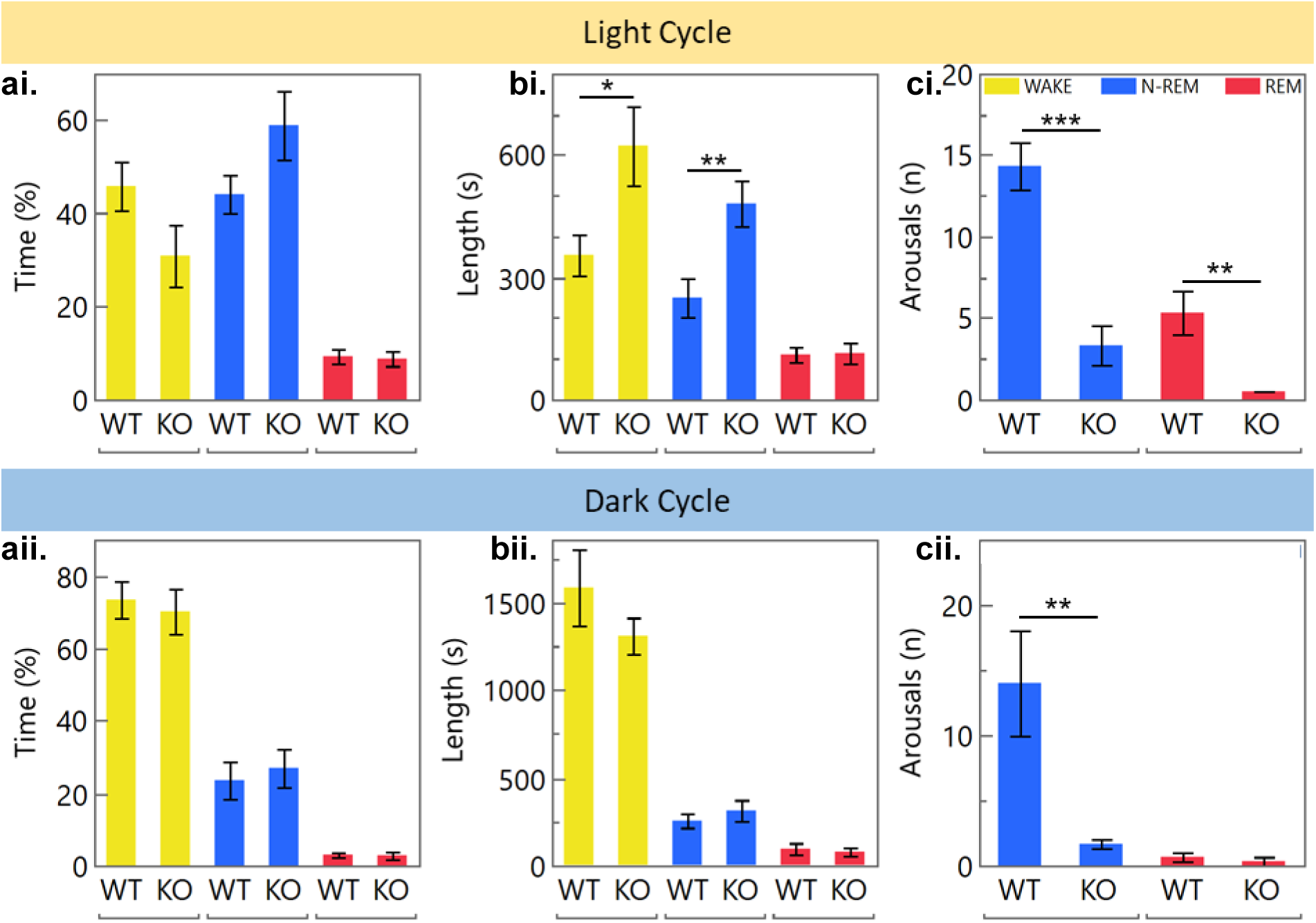
Sleep analysis of inducible IFT88 KO mice. (**a.**) Ablation of primary cilia in inducible IFT88 KOs does not alter the light or dark cycle sleep patterns of mice (**b.**) Average duration of wake and non-REM bouts was significantly increased (p=0.0454, 0.0179 respectively) in the light daytime cycle. (**c.**) Inducible IF88KO mice were found to sleep more soundly than their WT littermates as they were aroused to the awake state from non-REM (p=0.0024) and REM (p=0.0342) sleep significantly fewer times per sleep cycle than WT mice (Tukey’s HSD test), n=5 WT and 5 KOs.

### Inducible IFT88 KO mice have reduced EEG waveform activity

AC3 KO mice exhibit impaired neural activity, with a reduction in brain wave power similar to what has been observed in EEG recordings in human patients suffering from MDD ^38,39^. To characterize if IFT88 KO mice share this pathology the 24-hour EEG/EMG recordings were filtered to remove the DC offset as well as line current and then further processed in MATLAB to generate power spectrum density plots (PSDs; **Figure 3**). Inducible IFT88 KO mice had a broad reduction in brain power across the spectrum as well as a reduction in peak power during wake (p=0.0171) and non-REM (p=0.0341) mental states. More specifically, they lost approximately 47% of their power potential in the slow-wave delta-theta bands when in the awake state (p=.0013 for activity below 8Hz; **Figure 4**). When in non-REM and REM sleep, the slow wave power potential was not significantly disrupted, but instead there was a roughly 30% decrease in alpha and beta wave activity.

**Figure 3:**
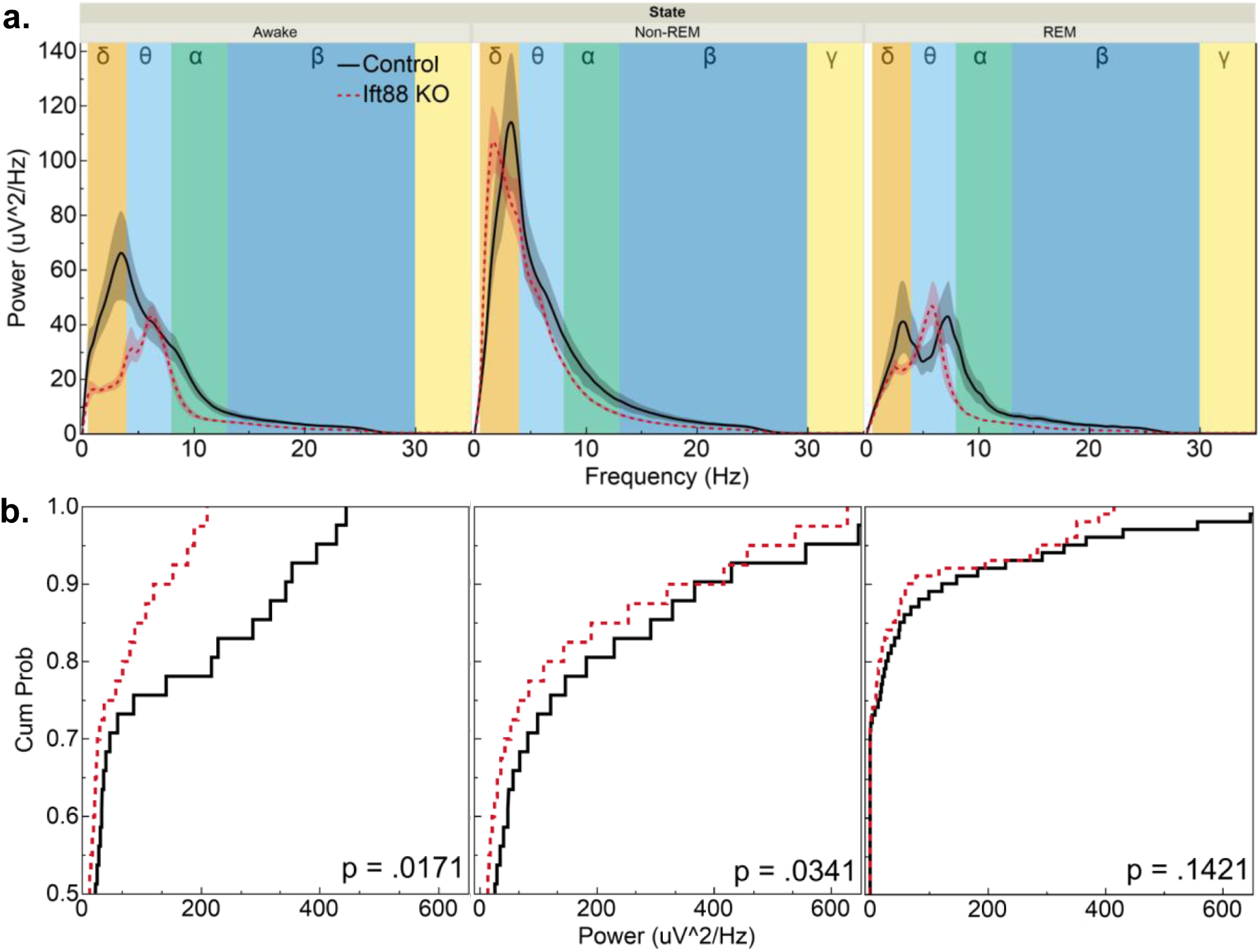
Inducible IFT88 KO mice have reduced EEG power over a broad spectrum. (**a.**) Power spectrum density plots (PSDs) of EEG waveform power broken down by frequency ranges reveals a shift in brainwave activity. Delta = 1 – 4 Hz, Theta = 4 – 8 Hz, Alpha = 8 – 13 Hz, Beta = 13 – 30 Hz, Gamma = 30 – 50 Hz. (**b.**) Cumulative distribution plots of the above PSDs show a shift in EEG waveform to a lower power. Overall EEG waveform activity was reduced in inducible IFT88 KOs in both the wake (p=0.017) and non-REM (p=0.034) states (Kolmogorov-Smirnov test).

**Figure 4:**
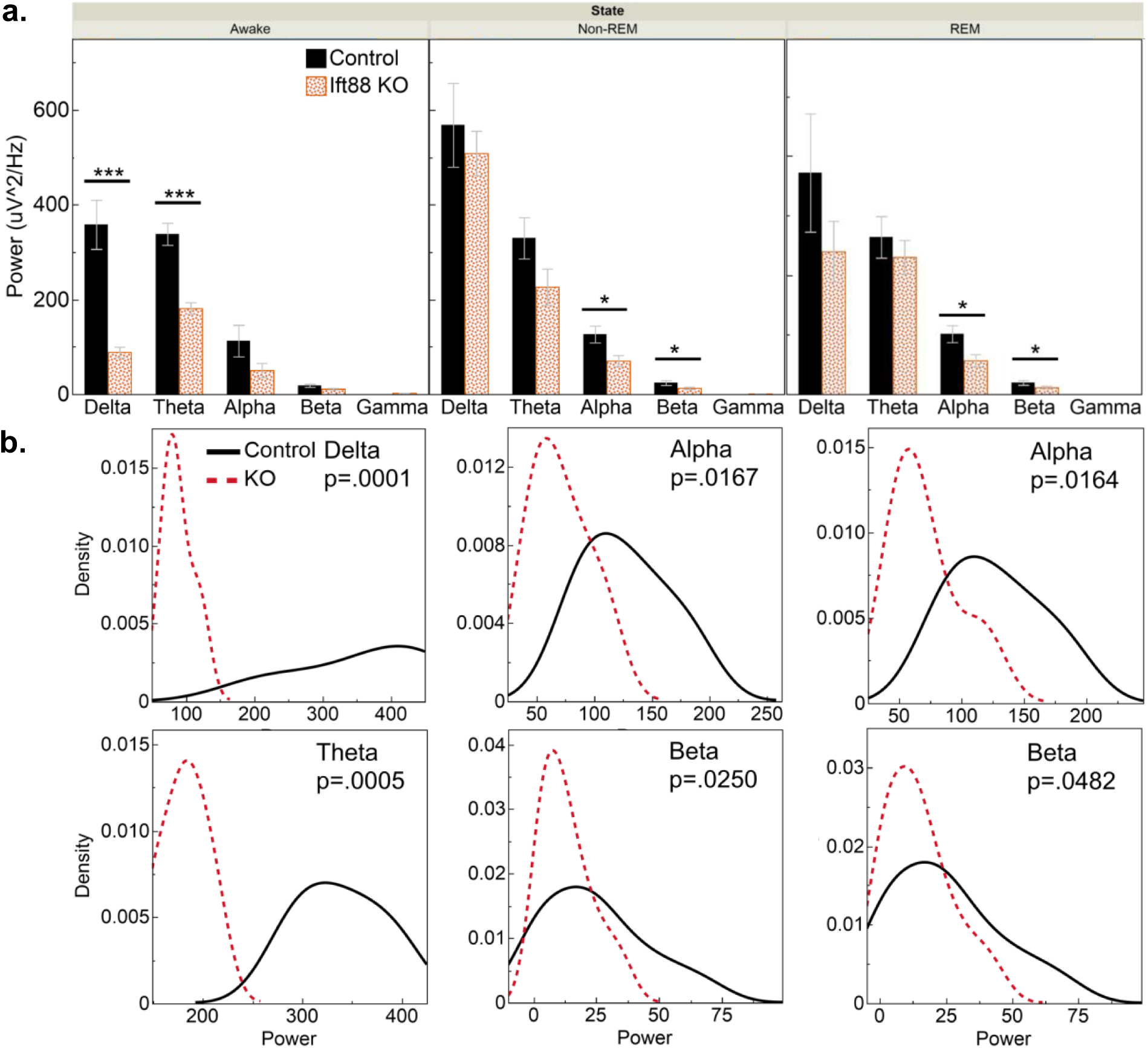
Ciliary ablation shifts EEG activity in a frequency dependent manner. (**a.**) Decomposition of PSDs reveals that during the awake state, low-frequency delta wave activity was reduced by 75.1% (p=0.0059) and theta wave power was attenuated by 52.8% in inducible IFT88 KOs (p=0.0013). This response was sleep-state dependent as slow-wave power was not attenuated in Non-REM and REM sleep. During sleep states only power of high frequency waveforms was impacted. During non-REM sleep alpha and beta wave power was reduced by 44.2% (p=0.0167) and 45.9% (p=0.0265). This remained consistent in REM sleep where alpha wave power was attenuated 43.9% (p= 0.0179) and beta wave power by 40.65% (p=0.0492) (Tukey’s HSD test). (**b.**) Density distribution plots of this data further illustrate how IFT88 ablation shifts peak brain power in these frequencies (Kolmogorov-Smirnov test).

### IFT88 ablation leads to attenuation of theta-gamma phase-amplitude coupling

It has been observed that different frequency bands of brain wave activity are not isolated and that their oscillations interact with each other through modulation (**Figure 5**). Multiple forms of this cross-frequency coupling (CFC) have been observed, however, phase-amplitude coupling (PAC) is the most studied form and is known to play a role in the integration of signals between different brain regions ^40,41^. In particular, slow-theta-to-gamma PAC in the human hippocampus has been found to support the formation of new episodic memories, and interactions between these bandwidths causes powerful and rapid action potential spiking leading to long-term potentiation at synapses ^42,43^. Slow-theta-to-gamma PAC has also been documented to be reduced or disrupted in patients that suffer from neurological diseases such as Parkinson’s disease, schizophrenia, and autism ^44–46^.

**Figure 5:**
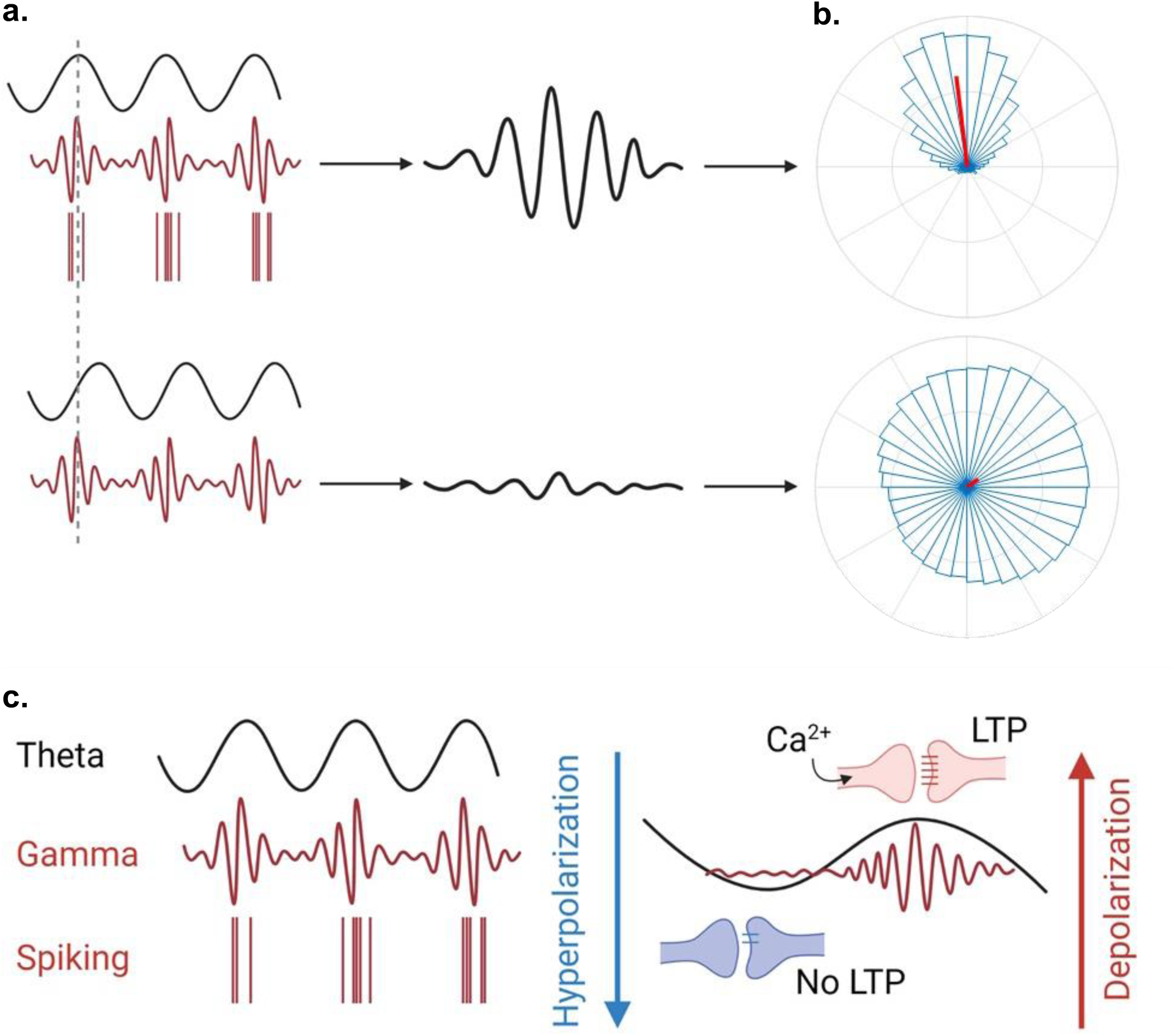
Phase-amplitude coupling. (**a.**) When the phases of different frequency waveforms harmonize, PAC is characterized by the formation of harmonic Morlet wavelets. (**b.**) In a polar histogram this can be quantified by a directional mean vector angle and mean vector length (red bar) which illustrates synchrony of the data set. When waveforms are not in phase they counteract each other, resulting in a lack of power and non-directional mean angle lengths and angles. (**c.**) Synchronization of slow wave (4-8 Hz) and high wave (30-100 Hz) waveforms initiate an influx of Ca^2+^ to depolarize the cell and trigger fast action potential spiking. This spiking leads to strengthening of synaptic connections and promotion of Long-term potentiation (LTP).

To quantify theta-gamma phase-amplitude coupling in IFT88 ablated mice, Canolty’s mean-vector length modulation index (MVL-MI) method was initially used (**Figure 6**). This method estimates PAC from a signal of length N, by combining phase and amplitude information to create a complex-valued signal: f ^ei(ffp)^ for every time point. This generates a table of vectors and if the resulting probability distribution function is non-uniform, it suggests coupling between the phase frequency (f_p_) and the amplitude frequency (f_a_), which can be quantified by taking the length of the average vector ^47^. First, for every experimental and control group PAC was calculated both with the pre-processed sample data and with a randomized version of the same dataset (to disrupt PAC) to calculate if PAC is present. Statistically significant theta-high gamma PAC was only found to be occurring in the awake and non-REM states of WT mice when compared to their matching pseudo-dataset (p<.0001). The Modulation Index (MI) was then compared between groups using Tukey’s HSD (note, a lower MI indicates stronger PAC). Using Canolty’s MVL-MI we found that IFT88 deficient mice had impaired theta-gamma PAC both when awake (p=0.0071) and when in non-REM sleep (p=0.0327). This is evident from the reduced mean vector length in both of these signals (p=0.0093 and p=0.0424) as well a 90° shift in the phase angle across all states (p=0.0366, 0.0383, 0.0415).

**Figure 6:**
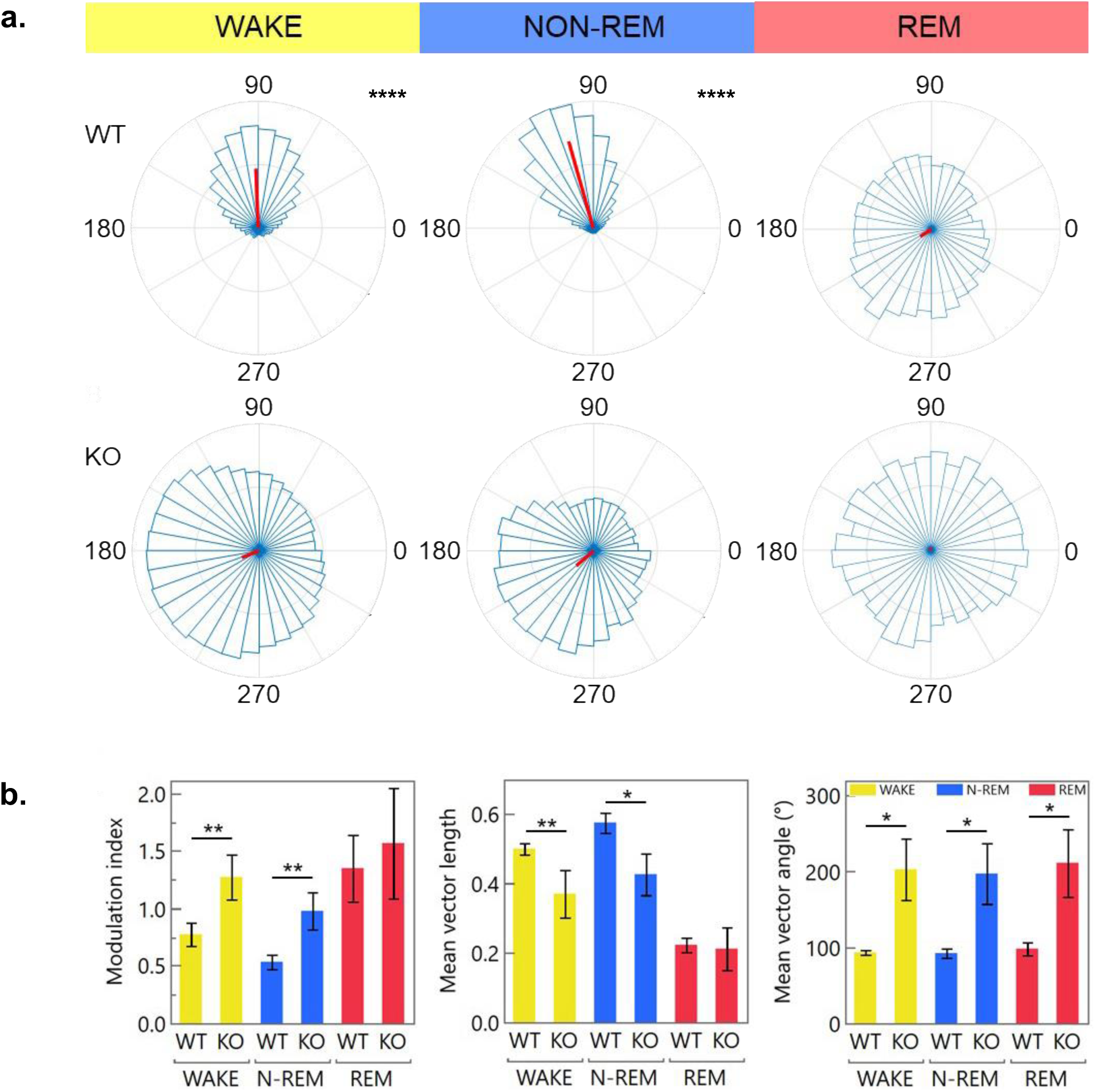
Inducible IFT88 KO mice have a reduced MVL-Canolty calculated PAC. (**a.**) Representative polar histograms of modulation index vectors calculated using the MVL-MI-Canolty method shows the formation of theta-gamma phase-amplitude coupling only in WT mice during wake and non-REM states (p<.0001). This coupling is weak or not present in inducible IFT88 KO mice (p>0.05). (**b.**) Using this algorithm a lower modulation index (MI) indicates stronger CFC. This reduction in PAC is quantified by a reduced mean vector length in wake (p=0.0071) and non-REM (p=0.0032) states as well as a 90° shift in their mean angle length vector in wake (p=0.0366), non-REM (p=0.0383), and REM (p=0.0415) states.

It should be noted, however, that MI-values from the MVL-MI-Canolty algorithm have been shown to partly reflect the power of f_a_ oscillations, rather than the coupling strength between f_p_ and f_a_. To further validate our conclusions, we applied a MI algorithm which includes a normalization factor that corresponds to the oscillatory power of f_a_ ^44^. This method has also been seen to be more resilient to noise, which is preferential for 2-lead EEG data that often has a lower signal to noise ratio than other electrophysiological recordings. Using this method, reductions in theta-gamma PAC strength during awake (p<0.01) and non-REM (p<0.05) activity were observed (**Figure 7**), consistent with our MVL-MI-Canolty based results. These findings suggest that the observed reduction in cross-frequency coupling is directly caused by the genetic ablation of IFT88 and is not due to electrical noise or variance.

**Figure 7:**
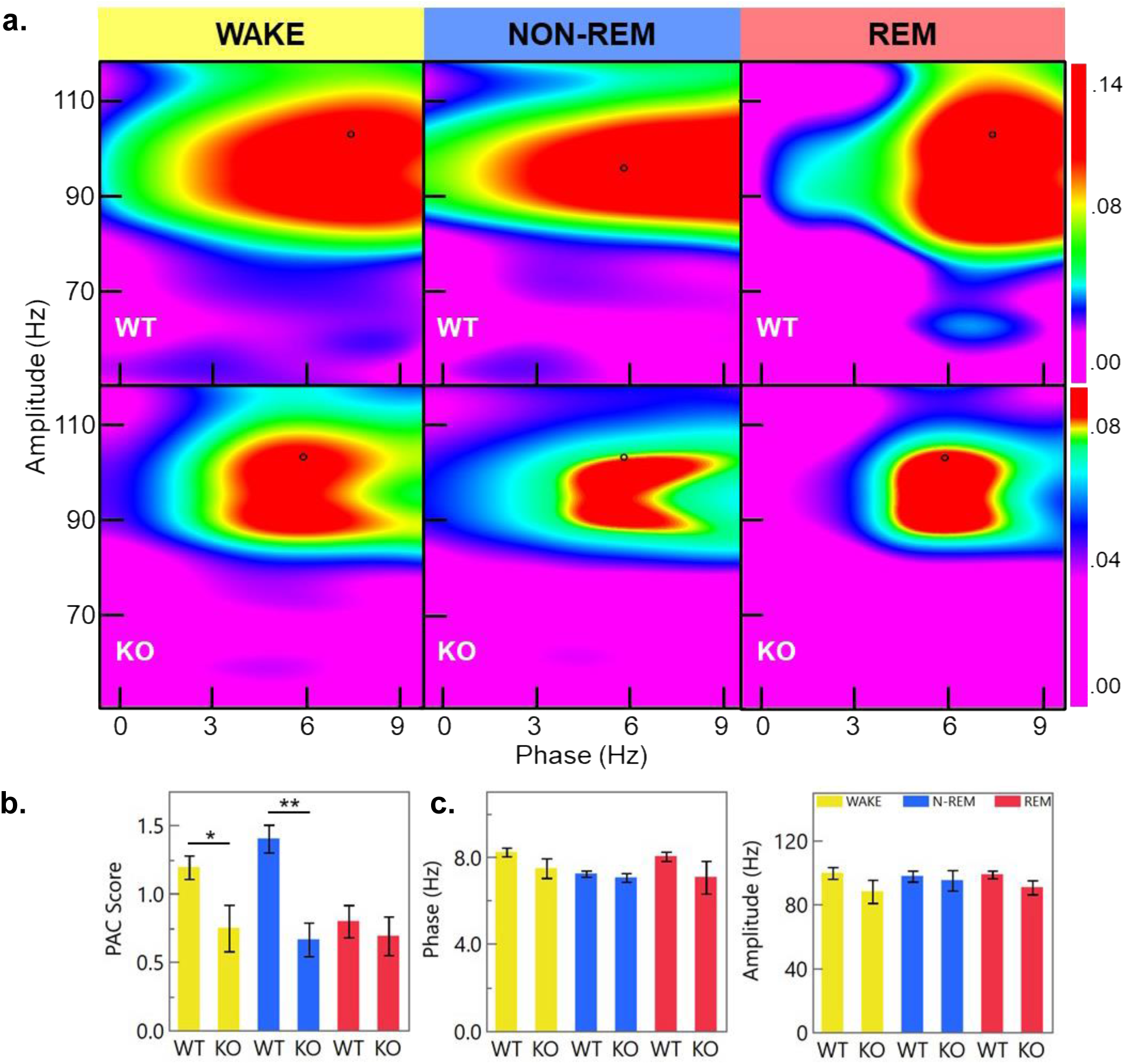
Theta-Gamma PAC is attenuated in inducible IFT88 KO mice. (**a.**) Representative heatmaps show phase-amplitude coupling between the theta-high gamma frequencies calculated using an adjusted MVL algorithm. Black cursors represent peak PAC for each condition. (**b.**) This algorithm corrects for oscillatory power in the signal used for PAC analysis to generate a PAC score value (a lower value indicates stronger PAC). Using this method inducible IFT88 KO mice still show a reduction in theta-high gamma cross frequency coupling during wake (p=0.0085) and non-REM (p=0.0372) states. (**c.**) Cross phase coupling is highest at the intersection of 8 Hz and 101 Hz in WT mice; this is not significantly changed in inducible IFT88 KO mice indicating that PAC of KO mice is being attenuated and not shifted.

## Discussion

Like AC3 KO depression model mice, inducible IFT88 KO mice were found to have altered sleep patterns, which is a common phenotype in human depression patients ^26,39^. The average sleep bout lengths of inducible IFT88 knockout mice were significantly longer than WT mice and they were aroused from sleep fewer times, indicating that inducible IFT88 KO mice are less likely to be stirred. This is possibly due to the state-dependent reductions in neural activity that were observed. Disrupted sleep patterns are also known to contribute to cognitive decline in Alzheimer’s patients where they are believed to interfere with natural amyloid-β and tau clearing processes ^48–50^.

Inducible IFT88 KO mice had no changes in their high frequency gamma and beta bandwidths but had a reduction in slow rhythm theta and delta waves. Interestingly, while awake the brain is predominantly active at higher frequencies, indicating that inducible IFT88 KO mice have impaired ability to generate low frequency subconscious processes while awake. The inverse of this phenomenon was observed in sleep states. In both REM and non-REM states inducible IFT88 KO mice had reduced high frequency activity in the beta and alpha frequencies, but no changes in the slow frequency theta and delta waves which are the predominate bandwidths during rest.

Our characterizations of an inducible IFT88 KO mouse model are similar but not identical to what has been seen in AC3 KO mice, highlighting it as a complementary mouse model for the study of primary cilia in neurological function ^26^. Differences are likely due to the near total extirpation of cilia seen in IFT88 KO mice, as well as the proteomic diversity of cilia. Apart from the core structural proteins required for all cilia, advanced protein tagging methods have demonstrated that the composition of these organelles can vary by over 70% between cell types^60–63^. Whereas AC3 KO limits the researcher to understanding the specific impacts caused by the loss of an adenylyl cyclase in neuronal cilia, IFT88 ablation allows for a broader starting point in which all possible ciliary signaling pathways are disrupted across various cell types. Our findings may also help to explain the memory and behavioral deficits that have been observed in other animal ciliary studies however further study is required to link specific ciliary pathways to cognition ^12–14,17–20,64,65^.

This is the first study to show that loss of primary cilia influences PAC, providing a new level of insight into how genetic mutations potentially influence the physical systems behind cognition and memory formation. Phase-amplitude coupling between the theta and gamma frequencies (TG-PAC) is an important electrophysiological process that aids in memory formation and is known to be disrupted in humans afflicted with mild cognitive impairment, Alzheimer’s, and Parkinson’s disease ^27,51,52^. Although the mechanisms behind the physiological propagation of TG-PAC remain unknown, recent single cell neurological recordings in human subjects revealed that PAC plays an important role in controlling interareal communication between the frontal and temporal lobes^53^. These findings that hippocampal PAC controls working memory supports the current hypothesis that PAC is a general neural mechanism for top-down control of complex bottom-up processes^54,55^. This implies that PAC likely also plays an important role in attention, decision making, speech comprehension, and memory retrieval^56–59^.

Consistent with our observations that cilia ablated mice exhibited signs of impaired associative and spatial memory, both of which are hippocampal dependent, we found that these mice also had severely depressed TG-PAC both while awake and in non-REM sleep. This finding not only supports the growing theorem that cilia play an essential role in adult neural function, but also provides novel evidence that ciliary signaling may be an important step in PAC formation.

Although it is not currently understood which ciliary signaling pathways contribute to brainwave activity, PAC, and hippocampal learning deficits, it is plausible that ciliary knock-out interferes with GPCR-mediated cAMP signaling within the cilium, ultimately causing synaptic dysfunction. IFT88 is a subunit of the IFT-B complex responsible for anterograde transport and elongation of the cilium ^68^. As ablation of this protein results in a lack of cilia formation, it will also disrupt numerous signaling pathways including the ciliary GPCRs SSTR3 and 5-HTR6, both of which have already been tied to cognition through genetic knock-out, pharmacological manipulation, and behavioral assays ^18–20^. It will additionally perturb ciliary AC3 signaling which has also been reported to influence contextual memory formation^69^. Adding further weight to the cAMP hypothesis are the findings that the primary cilium, despite its miniscule volume, is capable of generating cAMP signaling cascades independent of the cell body to influence adipogenesis and kidney cystogenesis ^70,71^. Previously this hypothesis would have been difficult to prove due to the complexities of measuring cAMP in live cells, but advancements in microscopy and the development of cilia specific cAMP biosensors will allow for further study of the primary cilium’s role in memory and neural function ^72^.

Classically, a role for cilia in cognition would have been considered unlikely, but our understanding of the primary cilium is constantly being redefined. Recent serial-section EM analysis revealed that each human cortical neuronal primary cilium forms connections with the processes and cell bodies of 40-60 other cells ^66^. A subset of these intersections develop synapse-like contacts, gap junctions, and other connections capable of propagating signals throughout the neural network ^66 67^. With this high level of neural integration, it is likely the primary cilium influences a host of functions yet uncovered.

Memory is one of the most complex yet fundamental traits of intelligent life, and when it is impaired by age or disease it can have devasting impacts both on the individual and those close to them. Our experiments and the work from other research groups previously cited are providing a growing body of evidence that primary cilia are necessary for proper neural function, not only in diseased patients but also in healthy individuals. Our findings that IFT88 ablation alters normal sleep patterning, EEG waveform activity, PAC, and impairs learning will help guide future research to elucidate the specific mechanisms that interlace memory and cognition with ciliary function.

## Methods and materials

### Animals and tamoxifen-induced ablation of IFT88 in mice

All animal-related husbandry, surgical procedures, and experimental protocols were approved by and conducted in accordance with the Institutional Animal Care and Use Committee (IACUC) of the University of New Hampshire (IACUC protocols #19040, #190502, #210801 and #211001) and meet or exceed ARRIVE guidelines. We confirm all experiments were performed in accordance with relevant guidelines and regulations. Mice were maintained on a 12-h light/dark cycle at 22°C and had access to food and water ad libitum. Mixed sex IFT88 flox/flox; UBC-Cre/ERT2 littermate mice of roughly two months of age were administered with tamoxifen dissolved in corn oil (0.2 mg/g body weight for consecutive seven days), or corn oil as controls, via oral gavage to induce CRE recombinase expression. Ift88 cilia KO mice exhibit normal acoustic and foot-shock responses. To verify that primary cilia were efficiently deleted in cilia KO mice, we assessed the efficiency of cilia ablation with immunostaining using AC3 antibody (1:4000, Cat# RPCA-ACIII, EnCor). More than 90% of primary cilia in the hippocampus were ablated using this inducible ablation protocol, consistent with our prior report^68^. For behavioral analyses, mice were handled by investigators for 3-5 days to allow them to adjust to the investigators before starting the experiments. After the treatment course mice were allowed to rest for six weeks to allow for full genetic silencing and turn-over of IFT88. Mice were monitored after injection and for 24 hours after their last injection for any signs of discomfort. All experiments used animals of mixed sexes. We didn’t note any significant differences in the results between male and female mice, so data were pooled. While our data show that both sexes are being affected by IFT88 deletion, we acknowledge that a larger sample size may reveal subtle differences.

### Trace Fear Conditioning Test

Trace fear conditioning was performed using inducible IFT88 KOs and controls 6-7 weeks after tamoxifen/vehicle administration. The mouse was placed in a fear conditioning chamber (VFC-008CT-LP) with a grid floor (VFC-005A, Med Associates Inc, Vermont). Trace fear conditioning was conducted as previously described^34^. After 5-10 minutes of exploration and acclimation, a neutral tone, which is the conditional-stimulus (CS) (3 kHz, 80 dB, 15 s) was delivered followed by a mild foot shock, which was the unconditional-stimulus (US) (1 second, 0.7 mA) 30 seconds later. The CS-US pairing repeated for seven cycles (10 s prior to tone, 15 s tone, 30 s delay, 1 s foot shock, 10 s post shock, at 190 s intervals) in which animals learned to associate CS with US and formed trace fear memory. After completion of the learning procedure, mice were then put into a box with bedding, food, and water ad libitum to rest for 2-3 hours before recall experiments. For recall experiments mice were placed into a novel environment distinct from the initial chamber to avoid contextual cues that would elicit contextual memory recall. Mice were subjected to five recall cycles with a tone stimuli and no foot shock (10 s prior to tone, 5 s with tone, 30 s delay, absence of foot shock, 10 s post shock, at 110 s intervals). The trace fear conditioning and recall experiments were videotaped using a high-resolution monochrome camera and freezing behaviors were analyzed offline. Videos were tracked using Noldus Ethovision XT (version 11.5) for real-time moving velocity, freezing, and locomotion. Freezing was defined as real-time velocity lower than 1 mm/s, as automatically measured by Ethovision XT. 10 pairs of inducible IFT88 KOs and controls (mixed sexes) were used for comparison in the trace fear conditioning test.

### Morris water maze test

The Morris water maze test was used to determine if inducible Ift88 KOs exhibit impaired spatial memory. A 150 cm-diameter circular pool was filled with 25 cm-deep water (23°C) made opaque with white tempera paint. Four shapes (square, circle, triangle and star) were labeled in the cardinal directions of the tank as reference points for the mice. A 10 cm-diameter circular platform was hidden 1.5 cm below the water surface in the target quarter. Three 120-second trials from different start points were conducted daily for 5 consecutive days. When a mouse failed to find the hidden platform within 120 seconds, it would be placed on the platform for 20 seconds. One day following training, mice were put back into the tank without the hidden platform for 120 seconds in the probe test (4 trials in the 6th day). The trials were videotaped (25 frames per second). Noldus Ethovision XT (Version 11.5) was used to track the latency to find the hidden platform, velocity, and duration in every quarter. Means of daily trials were compared between 10 pairs of (mixed sexes) inducible IFT88 KO and control mice.

### EEG/EMG headmount surgery

Surgery was performed as described previously ^69,70^ and followed the manufacturer’s instructions (Pinnacle Technology, Lawrence, KS, USA). Briefly, mice were anesthetized with either 1-2% isoflurane or an intraperitoneal injection of ketamine (130 mg/kg) /xylazine (8.8 mg/kg) cocktail and placed in a stereotaxic stage (Stoelting, Wood Dale, IL, USA). The scalp was then opened along the sagittal plane. Mice were secured in a stereotaxic device (Kopf Instruments, Tujunga, CA, USA). While under anesthesia, mice were prepared for surgery by hair removal, sterilization of skin with alcohol and 4% chlorhexidine. After exposing the skull surface, an EEG/EMG headmount (Pinnacle Technology Inc., Cat # 8201) was centered along the sagittal suture with the front edge 3.5 mm anterior to bregma, initially stabilized with VetBond. Headmount was then secured with four stainless steel screws (also functioned as EEG electrodes) and coated in acrylic for insulation. When positioned properly, all four screws (two anterior: AP 3.0 mm, ML ± 1.75 mm; two posterior: AP - 4.0 mm, ML ± 1.75 mm, relative to Bregma) sit on the cerebral region. Two EMG wires were inserted bilaterally into the trapezius muscles to monitor neck muscle activity. Skin was then sutured around the base of headmount. The headmount and surrounding screws and wires were then covered and sealed by dental cement.

### EEG/EMG recording and sleep state scoring

EEG/EMG recording were performed as previously described ^70^. Following surgery, the mice were given a week of recovery in their home cages before being individually tethered to EEG commutators within circular recording enclosures. They were then accustomed to the recording equipment in a free-moving state for two days. After acclimatization, continuous EEG/EMG signal collection was performed for a 24-hour period using Sirenia software from Pinnacle Technology (Lawrence, Kansas). The EEG/EMG signals underwent a 5,000-fold amplification and were sampled at 400 Hz. For classification into wakefulness, REM, and NREM sleep, EEG/EMG data were analyzed off-line. This process involved manual visual inspection supported by the cluster scoring method of Sleep Pro Version 1.3 (Pinnacle Technology), leveraging standard criteria that consider signal amplitude, frequency, and regularity. In 10-second epochs, EEG/EMG sleep/wake events were initially scored as wake (characterized by low-amplitude, high-frequency EEG, and high-amplitude EMG), NREM sleep (marked by high-voltage, low-frequency EEG, and low-amplitude EMG), or REM sleep (exhibiting low-voltage EEG predominantly in theta waves and EMG atonia). After scoring, segments representing different sleep states were exported as separate .edf files for MATLAB analysis.

### Data processing

Prior to analysis all signals were pre-processed in MATLAB R2020b using the open-source plugin Brainstorm v3 ^71^. Because phase-amplitude coupling and density distributions are time dependent, all files of wake, non-REM, and REM data had to be standardized to a consistent time period. This was done by concatenating all individual files for a specific sleep state of an individual into one file and then trimming to a standard length. For awake data this was 15,000 seconds per animal, non-REM was 10,000 seconds per animal, and REM was 2,000 seconds per animal. After concatenation each file was band-pass filtered between 0.5 and 250 Hz (Butterworth filter, low-pass order 4, high-pass order 3) and band-stop filtered (59.5–60.5 Hz; 119.5–120.5 Hz) to remove 60 Hz power-line contamination and its harmonics.

### Power spectrum densities

PSDs were calculated in Brainstorm using Welch’s method with a 1 second window time and 25% overlap. Frequency bandwidths were delta 0.5-4Hz, theta 4-8Hz, alpha 8-13Hz, beta 13-30Hz, gamma 30 - 200Hz. Significance was calculated using the Kolmogorov-Smirnov test.

### Phase amplitude coupling (PAC)

PAC was calculated in the MATLAB plugin EEGLAB v2021.0 ^72^. 3-10Hz was the phase frequency used for nesting against a high bandwidth of 50-120Hz. Modulation index was calculated using Canolty’s mean-vector length modulation index method ^47^.

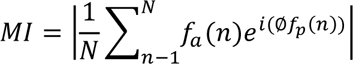

Phase-sorted amplitude histograms were compared using the Kolmogorov-Smirnov test with a Bonferroni-Holm correction and a 95% confidence interval. Initially to test if theta-high gamma PAC was present each sample dataset was compared to a pseudo-dataset where the initial data was randomized to disrupt and negate any possible PAC that was occurring. PAC was also calculated in Brainstorm using a modified modulation index algorithm that contains a normalization factor to correct for oscillatory power ^44^.

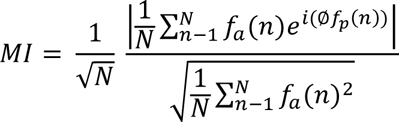

### Sample size and statistics

Ten pairs of inducible IFT88 KO and WT mice were used for behavioral analysis. Nine IFT88 inducible KO mice and 12 controls were subjected to isoflurane-induced anesthesia for isoelectric EEG recordings. For sleep-awake EEG waveform analysis five pairs of IFT88 KO and control mice were used. All groups contained mixed sexes and littermates were used to reduce genetic variation and behavior between treatment groups. After acquisition, labels were anonymized so the data could be scored blindly. Statistical comparisons were performed in JMP 15 Pro. First Shapiro-Wilk and Anderson-Darling goodness-of-fit tests were applied to evaluate normality of the data. Experiments with a normal distribution were compared using the Tukey-Kramer HSD while nonparametric results were compared using the Steel-Dwass test. For cumulative density plots of specific brain wave frequencies, the nonparametric Kolmogorov-Smirnov test was used to compare the distributions of brain wave activity.

## Author Contributions

XC and MS designed research; MS, YZ and LQ performed the research; MS, YZ and XC analyzed the data; XC and AMH provided resources; MS, YZ and XC prepared figure panels; MS drafted the manuscript; XC and AMH revised it.

## Acknowledgements

This study was enabled by NIH Grants K01AG054729, P20GM113131, R15MH126317 and R15MH125305, COLE Neuroscience Research Awards, and UNH CoRE PRP awards to XC, and VA-ORD 1I01BX005124 to AMH. MS and YZ were supported by Summer TA Research Fellowships (STAF) and a Dissertation Year Fellowship (DYF) of UNH Graduate School. We thank the University Instrumentation Center for A1R HD confocal imaging service.

## Data availability

All original raw EEG recordings have been preserved and are available as Pinnacle Technology filetypes. Additionally, processed files that were generated in MATLAB are available upon request. All data values used in statistical analysis are available in .csv filetypes if requested. To request any data please reach out to the corresponding author at.

## Ethics Declaration

All animal-related husbandry, surgical procedures, and experimental protocols were approved by and conducted in accordance with the Institutional Animal Care and Use Committee (IACUC) of the University of New Hampshire and meet or exceed ARRIVE guidelines. IACUC protocols #190403, #190502, #210801 and #211001. We confirm that all experiments were performed in accordance with relevant guidelines and regulations.

## Supplementary Figure

**Figure S1:**
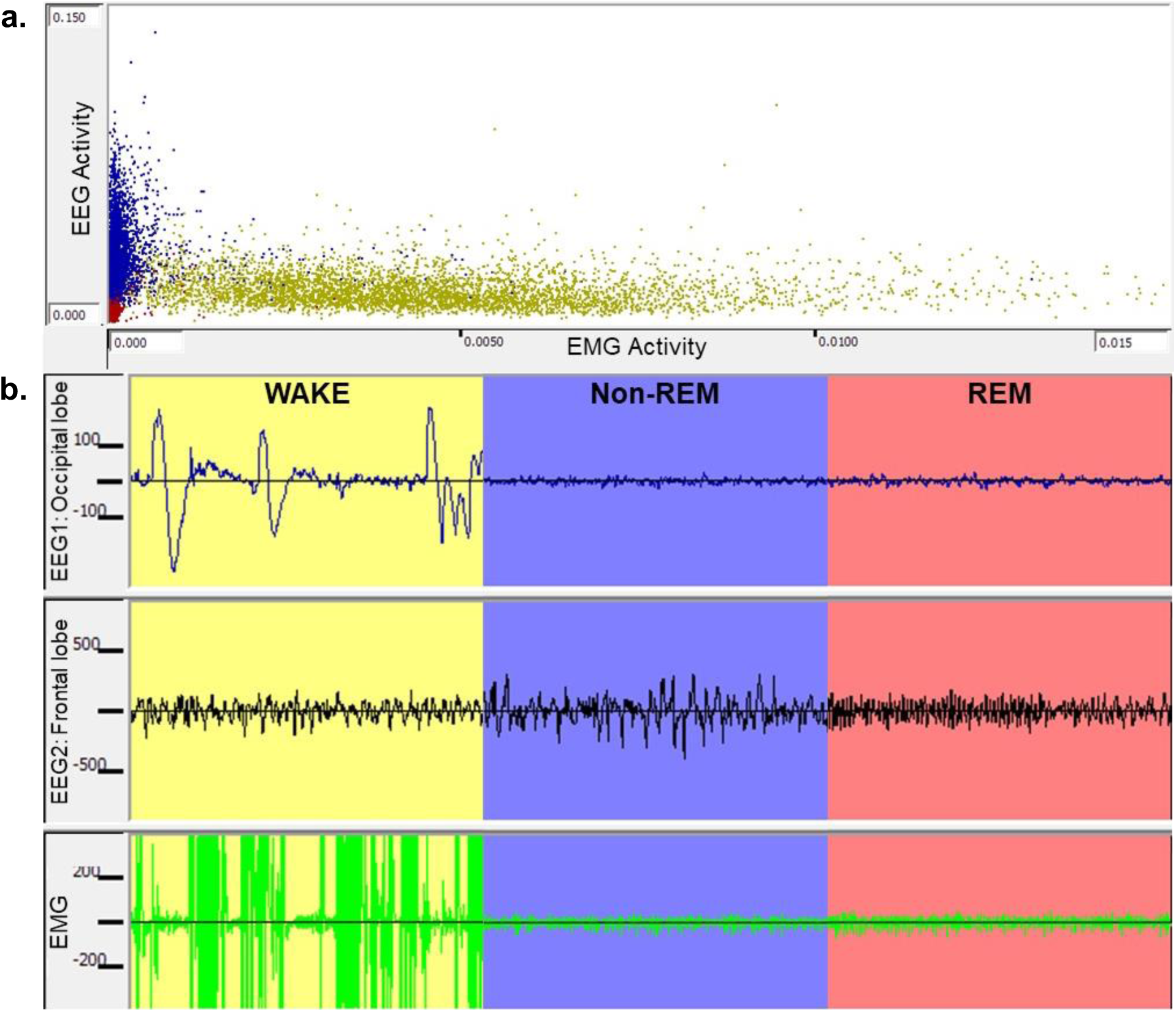
Manual scoring of mouse EEG/EMG electrophysiological recordings. (**a.**) Example of how 24-hour recordings were cluster-scored in Sirenia Sleep 1.0.3 by comparing EEG to EMG signal strength to identify clusters of Wake, Non-REM, and REM data. (**b.**) The recordings were manually curated in 10 second epochs to correct cluster scoring. The awake state is characterized by activity in the occipital lobe (EEG1), high frequency-low amplitude activity in the frontal lobe (EEG2), and high levels of EMG activity. Asleep mice lack EMG and occipital lobe activity. The non-REM state is identified by low frequency/high amplitude brain waveforms and REM sleep is differentiated by high frequency-low amplitude EEG activity, similar to what is seen while awake.

